# NeuroCarta: An Automated and Quantitative Approach to Mapping Cellular Networks in the Mouse Brain

**DOI:** 10.1101/2025.03.25.645187

**Authors:** Tido Bergmans, Tansu Celikel

## Abstract

Understanding the structural organization of the brain is essential for deciphering how complex functions emerge from neural circuits. The Allen Mouse Brain Connectivity Atlas (AMBCA) has revolutionized our ability to quantify anatomical connectivity at a mesoscale resolution, bridging the gap between microscopic cellular interactions and macroscopic network organization. To leverage AMBCA for automated network construction and analysis, here we introduce NeuroCarta, an open-source MATLAB toolbox designed to extract, process, and analyze brain-wide connectivity networks. NeuroCarta generates directed and weighted connectivity graphs, computes key network metrics, and visualizes topological features of brain circuits. As an application example, using NeuroCarta on viral tracer data from the AMBCA, we demonstrate that the mouse brain exhibits a densely connected architecture, with a degree of separation of approximately four synapses, suggesting an optimized balance between local specialization and global integration. We identify attractor nodes that may serve as key convergence points in brain-wide neural computations and show that NeuroCarta facilitates comparative network analyses, revealing regional variations in projection patterns. While the toolbox is currently constrained by the resolution and coverage of the AMBCA dataset, it provides a scalable and customizable framework for investigating brain network topology, interregional communication, and anatomical constraints on mesoscale circuit organization.

## Introduction

Despite the fundamental simplicity of individual neurons, their collective organization into large-scale circuits gives rise to sophisticated behaviors and computations (Azarfar, Calcini, et al., 2018; Carandini, 2012; Gu et al., 2015; Huang et al., 2022; Sandamirskaya et al., 2022). Understanding the brain’s structural organization is thus crucial for deciphering the principles underlying neural information processing.

At the mesoscale level—an intermediate resolution bridging microscopic cellular connections and macroscopic regional interactions—mapping anatomical connectivity reveals fundamental principles of neural information processing (Hilgetag & Hütt, 2014; Huang et al., 2022; Scheenen & Celikel, 2015; Senk et al., 2022), sensorimotor integration (Heckman et al., 2017; Oh et al., 2014; Rault et al., 2024), and cognitive function (Paquola et al., 2025; Suárez et al., 2020). The Allen Mouse Brain Connectivity Atlas (AMBCA) has emerged as a landmark resource to quantify anatomical connectivity across the entire mouse brain (Oh et al., 2014). By providing a standardized three-dimensional reference framework (Kuan et al., 2015; Wang et al., 2020) based on viral tracer mapping, the AMBCA enables researchers to explore global and local connectivity patterns, advancing our understanding of brain network architecture. Moreover, the extensive dataset of projection mappings based on targeted neuronal populations provided by AMBCA augments previous gene expression datasets, allowing for a deeper understanding of the biological mechanisms underlying connectivity formation and functionality (Fakhry & Ji, 2015; Takata et al., 2021). This integration of multidimensional data—gene expression coupled with anatomical mapping—facilitates the construction of a comprehensive view of mesoscale brain networks, bridging structural and functional analyses (Grandjean et al., 2017).

Several studies have successfully leveraged the neuroanatomical organization obtained from the AMBCA to quantify the connectivity of specific brain regions. In the original publication introducing the database, Oh et al. (2014) demonstrated that cortico-cortical connections broadly follow a lognormal distribution of strengths, with some connections stronger than a simple spatial dependence model predicted. They also revealed that functional network organization mirrors the underlying structural connectivity, particularly the distinction between ipsilateral and contralateral projections. Subsequent work has built upon this, revealing hierarchical organization within cortical networks, and identifying modular structures that reflect functional specialization (Knox et al., 2018).

Going beyond cortical connectivity, the AMBCA has been crucial for detailing the projections from cortex to various subcortical structures. Oh et al. (2014) provided an initial overview, highlighting the topographic organization of projections to the striatum and thalamus. The AMBCA showed that the striatum can be segregated based on differential resting-state fMRI connectivity patterns which mirror the monosynaptic connectivity with the isocortex; the functional connectivity between these cortico-subcortical regions can emerge via monosynaptic and polysynaptic pathways (Grandjean et al., 2017). Further studies delved into specific pathways, such as the cortico-pontine projections (Øvsthus et al., 2024) and the somatosensory and motor cortices, revealing details of their modular organization (Heckman et al., 2017; Rault et al., 2024). Additionally, advancements in imaging techniques have positioned the Allen Brain Atlas as a crucial reference point for cross-modal comparisons. Takata et al. (2021) demonstrated the feasibility of integrating imaging modalities such as MRI, DTI, and fMRI with the AMBCA dataset, allowing for multi-resolution analysis of brain networks. These computational approaches promote a more comprehensive understanding of how anatomical structure supports neural function.

As mesoscale connectivity mapping becomes increasingly central to neuroscience, automated computational tools are required to extract, analyze, and interpret the vast amounts of data generated by the AMBCA. Several network analysis approaches have been developed, e.g. (Friedmann et al., 2020; Knox et al., 2018), and used to study connectivity across the mouse brain as described above, but existing methods often require specialized programming expertise, lack comprehensive analytical pipelines, or focus on specific network properties rather than providing an integrated solution. To address these limitations, we introduce NeuroCarta, an open-source MATLAB-based toolbox designed to facilitate automated, large-scale network construction and analysis using AMBCA data. NeuroCarta enables researchers to construct weighted and directed network representations of the mouse brain, facilitating the investigation of global and local connectivity properties. The toolbox supports automated data extraction, connectivity matrix generation, and advanced network analysis, quantifying key network properties such as degree of separation, clustering, hub connectivity, and interhemispheric projections. As application examples we analyze the structural connectivity of the mouse brain using NeuroCarta, revealing that the network is densely connected, with a degree of separation of approximately four synapses, indicative of high computational efficiency. Additionally, we identify key attractor nodes with significantly higher input-to-output ratios, which may serve as critical hubs for information integration and relay processing. Comparative analyses also highlight sex-specific differences in connectivity, particularly within sensorimotor circuits, further exemplifying how NeuroCarta provides a powerful tool for exploring the anatomical foundations of information flow in the mouse brain.

## Methods

NeuroCarta (https://github.com/DepartmentofNeurophysiology/Neurocarta/) is a plug-and-play, open-source MATLAB toolbox facilitating the construction and analysis of mouse brain neural networks using data sourced from the Allen Mouse Brain Connectivity Atlas (AMBCA). Its data pipeline encompasses data download and import, network compilation, and network analysis and visualization tools (**Figure 1**). The default workflow generates a mesoscale, bilateral connectome of the mouse brain, but users can readily customize the network creation process through user input and metadata at various pipeline stages. This flexibility allows for the generation of tailored networks, such as those focused on specific connection types (e.g., excitatory only) or defined brain circuits; see **Supplemental Figure 1** for an example of the output.

**Figure 1.**
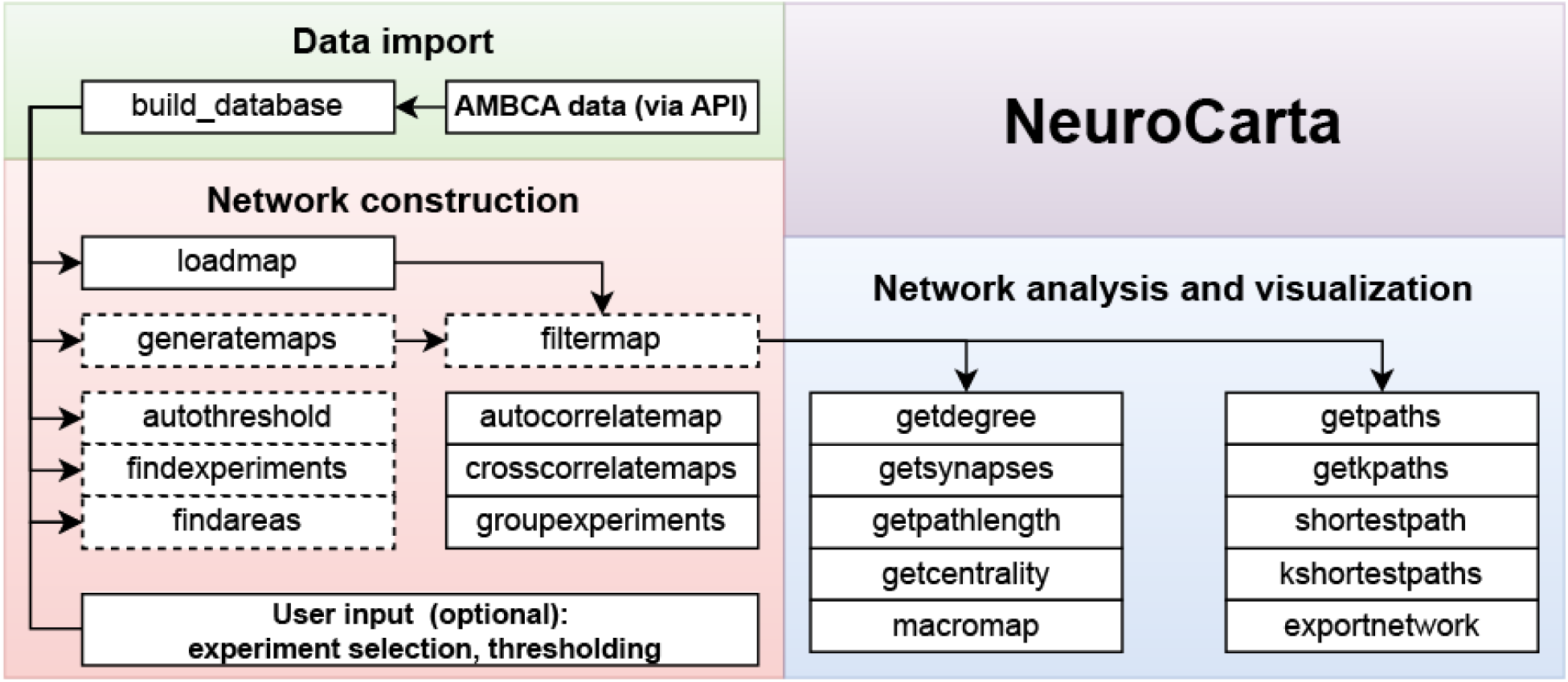
Overview of the toolbox functionality and workflow.

### Data import

NeuroCarta’s *build_database* function leverages the AMBCA application programming interface (API) to import experimental data. By default, this function downloads the entirety of the AMBCA dataset, which currently comprises 2918 brain imaging experiments. Alternatively, users can provide a curated list of experiments obtained, for example, through targeted searches on the AMBCA website, to constrain the data import to specific experimental subsets. The download process is designed to be robust; it can be interrupted and resumed later, allowing for incremental data acquisition.

In addition to the core experimental data, *build_database* retrieves associated metadata, encompassing (but not limited to) transgenic mouse lines, mouse strains, sex, and injection volume. Furthermore, metadata regarding the AMBCA reference atlas, including stereotaxic coordinates of brain structures and their hierarchical relationships, are also imported.

Each AMBCA experiment represents a brain imaging procedure involving the injection of a fluorescent protein-expressing, anterograde viral tracer into a precisely defined location within a genetically modified mouse brain. In a subset of experiments, Cre/loxP mediated gene recombination ensures the targeting of genetically defined cells. Within these cells, the fluorescent protein distributes throughout the axonal arbor but does not cross synapses. Image segmentation is performed on the acquired fluorescence images, quantifying the relative axonal density originating from the injection site and projecting to other regions of the bilateral brain. Although four distinct measures are available (projection density, projection intensity, projection energy, and projection volume), NeuroCarta’s default network construction utilizes projection density. The user can select any of the other three to reconstruct the networks.

Downloaded experiment data, stored in JSON format, undergoes a series of processing steps. First, the injection hemisphere is computationally determined. Experiments failing to meet the following criteria are discarded to improve accuracy: (1) a greater number of structures with the *is_injection* property set to true; (2) a higher total sum of projection densities; and (3) the presence of the structure exhibiting the highest projection density—all within the same hemisphere. The hemisphere meeting these criteria is designated as ipsilateral (relative to the injection site), and the opposite hemisphere as contralateral.

Subsequently, the injection structure is identified as the structure within the ipsilateral hemisphere exhibiting the maximum projection density. This determination is restricted to a predefined list of 302 non-overlapping brain structures covering the entire brain, stored in *nodelist.mat*. This list is derived from the AMBCA reference atlas, but users can substitute a custom list, enabling the construction of networks at different resolutions.

Finally, the projection data within each experiment are normalized with respect to the designated injection site. The injection site is assigned a projection density of 1.0, and all other projection densities within that experiment are scaled to the interval [0, 1]. The processed data are then stored in the MAT file format.

Imported data are accessible for individual-experiment level refinement and inspection. Functions such as *findarea* and *findexperiments* allow users to identify regions and experiments meeting specific criteria. *autothreshold* and *filtermap* enable noise reduction. The functions *autocorrelatemap* and *crosscorrelatemaps* provide extensive visualizations of statistical properties from a single or pair of experiments (see Supplemental Figure 1).

### Network construction

NeuroCarta’s network construction is primarily facilitated by the *loadmap* function. This function compiles data from individual experiments, which represent monosynaptic axonal projections (defaulting to projection density), into a comprehensive, polysynaptic network representation. The user can specify a subset of experiments for inclusion, or, by default, *loadmap* processes all available experiments within the imported dataset.

The core of network construction involves populating a connectivity matrix (adjacency matrix). This matrix is structured such that each row corresponds to a source node (brain region or voxel, depending on the resolution), and each column corresponds to a target node. Crucially, each AMBCA experiment provides data for a single row of this matrix, representing the outgoing connectivity from the experiment’s identified injection site.

If multiple experiments share the same injection site (as determined by the *nodelist.mat* or a user-provided equivalent), the corresponding rows in the connectivity matrix are averaged to produce a single, representative row. Nodes not designated as injection sites in loaded experiments are excluded from the resulting network. This is a form of source-based parcellation. The resultant connectivity matrix is inherently bilateral, reflecting the organization of the AMBCA data. It is sized N x 2N, where N is the number of included nodes. The first N columns represent ipsilateral connectivity (targets on the same side of the brain as the injection site), and the subsequent N columns represent contralateral connectivity (targets on the opposite hemisphere). Downstream analysis functions within NeuroCarta are designed to accept both unilateral (N x N) and bilateral (N x 2N) matrices.

Additional functions provide flexibility in network generation. *generate_maps* facilitates the batch creation of multiple networks by iterating over a user-defined set of parameters, generating a distinct network for each parameter combination. The *groupexperiments* function allows for constructing a single row of the adjacency matrix, representing the outgoing connections from grouped experiments.

### Network analysis and visualization

The NeuroCarta toolbox provides a suite of functions for analyzing and visualizing the constructed networks. These functions operate on the connectivity matrix (described in the “Network Construction” section) and provide node- and network-level metrics.

**Node-Level Metrics**

- **Degree:** The fundamental node-level metric is the degree computed by the *getdegree* function. Because NeuroCarta constructs directed networks, each node has an in- and out-degree.

- In-degree: The sum of all incoming connection weights (projection densities) to a given node.
- Out-degree: The sum of all outgoing connection weights (projection densities) from a given node.

**Pathways and Distances**:

NeuroCarta focuses on analyzing pathways and distances within the network. A key concept is the edge weight, which, in contrast to projection density, represents a distance between connected nodes. Edge weight is defined as the inverse of the projection density:

- **Edge Weight:** edge weight(i, j) = 1 / projection density(i, j) An optional multiplicative factor can be incorporated to represent the “cost” or “weight” associated with crossing a synapse.
- **Path:** A sequence of connected edges between a source node and a target node.
- **Path Length:** The sum of the edge weights along a given path: Path length = Σ edge weight(i, j) for all edges (i, j) in the path. The *getpathlength* function provides this to the user.
- **Weighted Distance (Shortest Path):** The minimum path length between two nodes, calculated using Dijkstra’s algorithm. The *shortestpath* and *getpaths* functions implement this algorithm.
- **k-Shortest Paths:** An extension of Dijkstra’s algorithm, implemented in *kshortestpaths* and *getkpaths*, computes not only the shortest path but also the k next-shortest paths between two nodes.
- **Betweenness Centrality:** Computed by the *getcentrality* function. Represents the fraction of all shortest paths within the network that pass through a given node. This provides a centrality measure for each node.
- **Relative Density** is used for comparative analysis of networks, used in this research to quantify sex-specific networks: Relative density = (Density_male_ - Density_female_)/(Density_male_ + Density_female_)

**Binary Network Analysis**:

Neurocarta also provides functionality for analyzing unweighted, binary networks.

- **Binarization:** A weighted network can be converted to a binary network by applying a threshold to the edge weights. Edges with weights below the threshold are set to 0 (representing absence of connection), and those above the threshold are set to 1 (representing presence of connection).
- **Degree of Separation (DOS):** In a binary network, the shortest path length, computed via Dijkstra’s algorithm, directly corresponds to the number of edges (and therefore, the minimum number of synapses) separating two nodes. The *getsynapses* function calculates this “degree of separation.”
- **Weighted DOS:** Number of synapses crossed along the shortest path between the two nodes in the weighted network.

**Network Export and Visualization**:

- **exportnetwork:** This function exports the network data in the .gexf (Graph Exchange XML Format) file format. This format is compatible with popular network visualization and analysis software such as Gephi (Bastian et al., 2009). This allows users to leverage external tools for advanced visualization and analysis.
- **Fruchterman-Reingold**: Within Gephi, the Fruchterman-Reingold layout algorithm (Fruchterman & Reingold, 1991) is recommended for visualizing network structure, including identifying clusters.
- **macromap:** This Neurocarta function generates a condensed version of the network by averaging node properties (e.g., connectivity, degree) within larger, user-defined brain areas. This provides a higher-level view of network organization.

## Results

NeuroCarta can be used to systematically study the mesoscale connectome of the mouse brain. Below, through quantitative network analysis, we exemplify its use, focusing on emergent properties such as the dominant ipsilateral connectivity, the heterogeneous input/output profiles of individual nodes, and the spatial dependence of connection strengths. We further showcase the toolbox’s capacity by dissecting sex-specific network architectures and the organization of the sensorimotor circuits within this comprehensive dataset.

### Connectivity Patterns and Network Structure

Using the NeuroCarta toolbox, we constructed a directed, weighted network of the mouse brain based on projection density data from the Allen Mouse Brain Connectivity Atlas (AMBCA; (Oh et al., 2014)). The resulting mesoscale connectome consists of 276 nodes per hemisphere (brain regions) with over 140,000 directed edges prior to any thresholding. Edge weights (projection densities) were normalized to the range [0, 1], and a bilateral adjacency matrix of size 276x552 was obtained (Figure 2A). The network was exported and in Gephi a network layout was generated using the Fruchterman-Reingold algorithm (Supplemental Figure 3).

**Figure 2.**
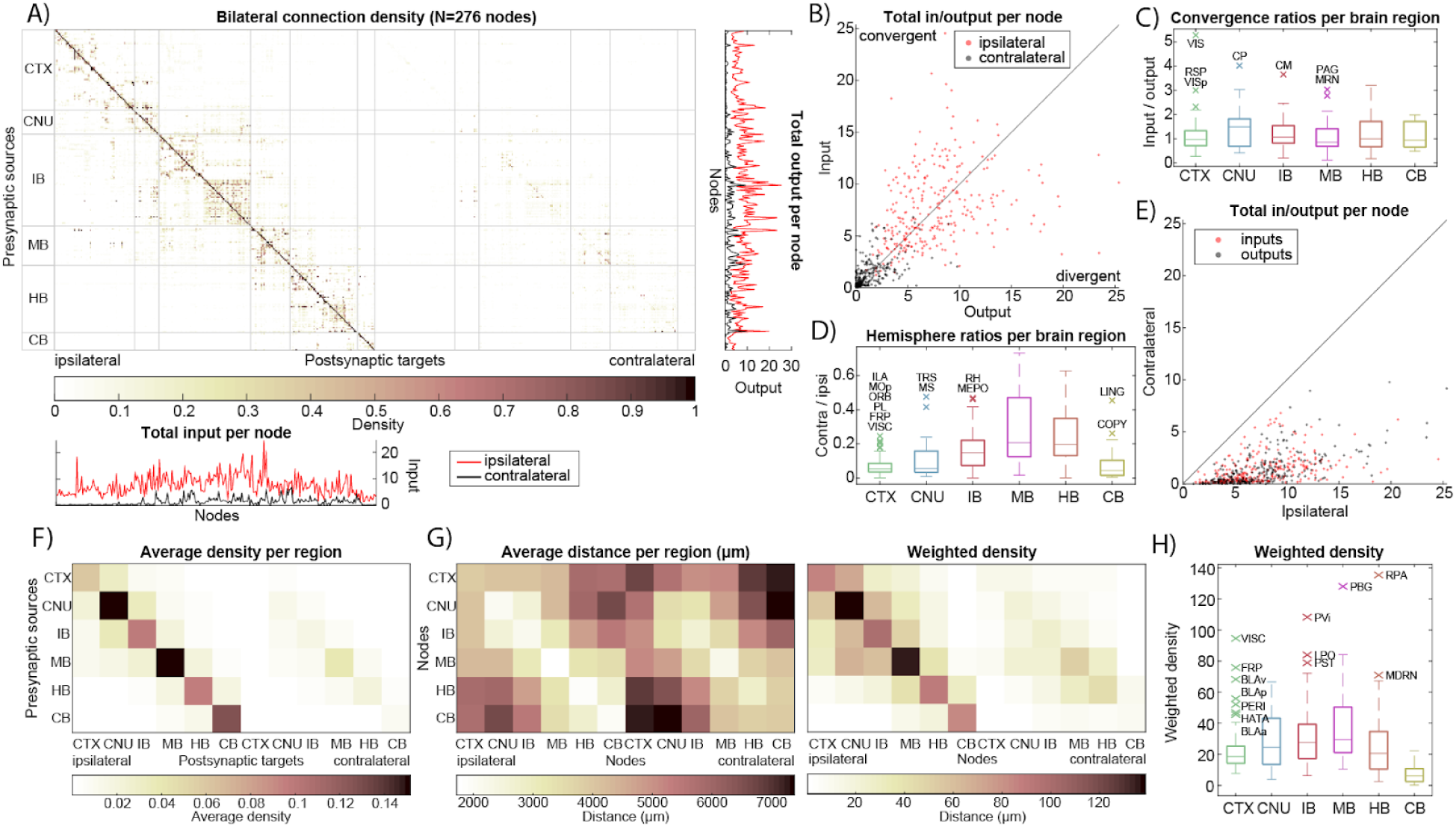
Bilateral connectivity of the mouse brain: adjacency matrix and nodal properties. **A)** Adjacency matrix of the 276-node bilateral brain network (size: 276x552) based on projection density. Line plots on the bottom and right, respectively show the total input vs output (i.e. sum over columns resp. rows of the adjacency matrix) per node. Data is shown separately for ipsilateral (red) and contralateral (black) projections. **B)** The total input and output of nodes for ipsilateral and contralateral projections reveal convergent and divergent nodes. **C)** Node convergence (input/output) ratio, shown separately per larger brain region. Outliers are denoted with their node acronyms taken from AMBCA (see supplemental table 1). **D)** The total node input and output per hemisphere reveal a preference for ipsilateral connectivity. **E)** Node hemisphere ratios, shown separately per larger brain region. **F)** Projection density-based adjacency matrix condensed by averaging nodes per larger brain region. **G)** Left: the distance in micrometers between nodes is averaged per larger brain region. Right: weighted density (density*distance) per larger brain region. **H)** Node weighted density, shown separately per larger brain region.

In the mouse brain, ipsilateral connections dominate the network: 84.7% of all connections are confined to the same hemisphere, i.e. most projections from a given region terminate in regions of the same hemisphere. There is also a strong autoconnectivity effect, wherein 70.1% of those ipsilateral connections occur within the same higher-level brain division (e.g., cortex-to-cortex, thalamus-to-thalamus). This intra-division bias is evident when we aggregate the connectivity matrix by major brain regions (Figure 2F), which shows that within-region connectivity far exceeds inter-region connectivity. Interestingly, while ipsilateral links carry higher weights on average, contralateral connections are more numerous: when projection densities to each hemisphere are normalized separately, one can see many low-density contralateral links that do not appear ipsilaterally (Figure 3A,B). In fact, in the unfiltered network, about 92% of all possible region-to-region connections exist (mostly weak projections). Thus, although cross-hemisphere projections tend to be weaker than same-side ones, they span a wider variety of region pairs, contributing to the dense, near-complete connectivity of the overall network.

**Figure 3.**
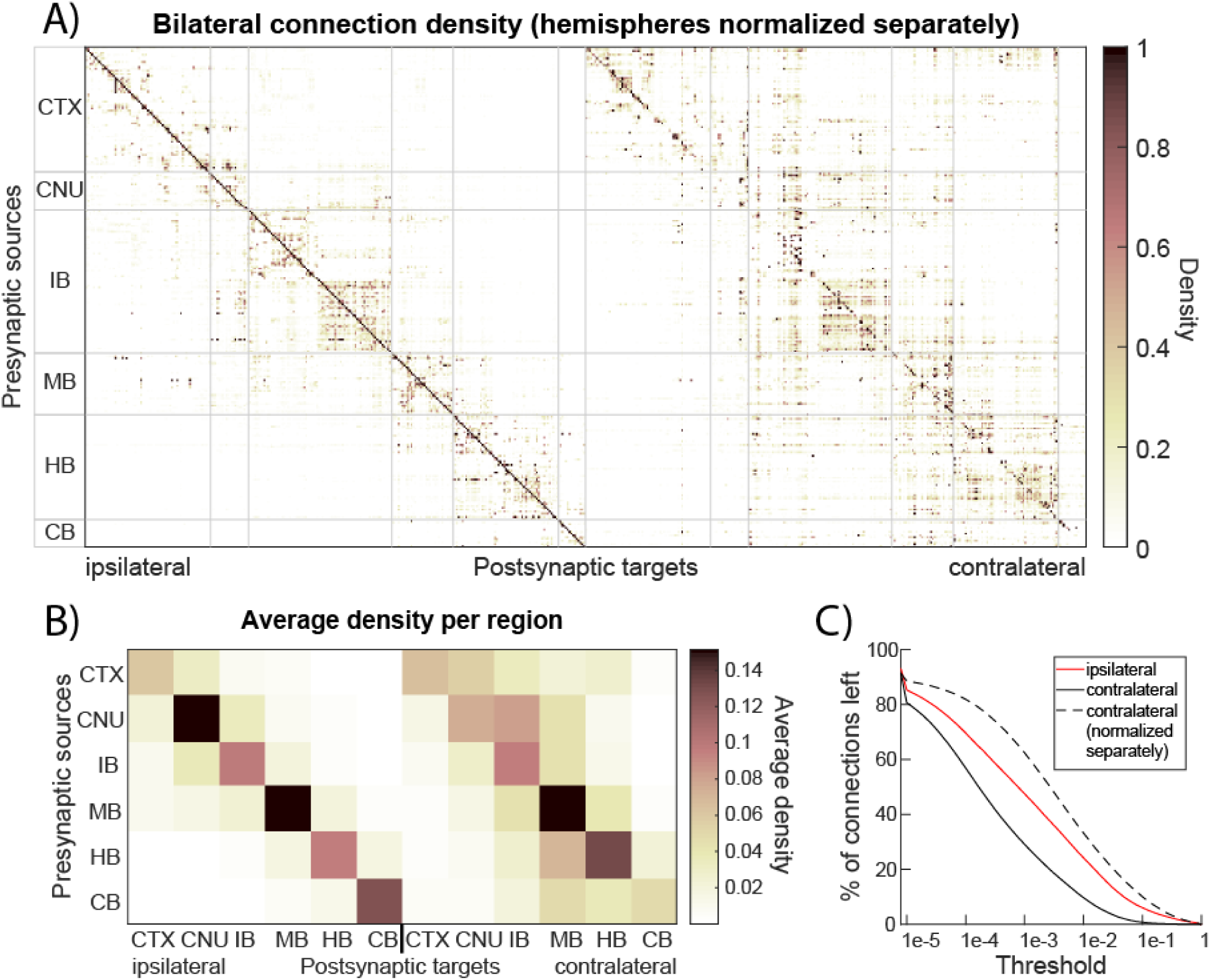
Bilateral connectivity of the mouse brain after normalizing connectivity separately in each hemisphere. **A)** Projection density-based adjacency matrix with projections normalized separately for both hemispheres. **B)** Adjacency matrix from A) condensed by averaging projections per larger brain region. **C)** Effect of thresholding on resulting number of ipsi- and contralateral projections.

At the node level, there is substantial variability in the balance of inputs and outputs across different brain areas. Some nodes act as convergence hubs, receiving disproportionately more input than they send out, while others are divergence hubs with strong outputs relative to their inputs. This is illustrated in Figure 2B, which plots the total input vs. output for each node and reveals nodes above the unity line (convergent, net receivers) and below it (divergent, net senders). A subset of regions show significantly high input/output ratios (marked as outliers in Figure 2C), indicating they integrate information from many sources. Conversely, a few regions have much higher outbound connectivity than inbound. In particular, several nodes exhibit a pronounced contralateral projection bias: they send a large fraction of their total output to the opposite hemisphere. Several areas project up to ∼50% of their outputs contralaterally, far above the norm. These contralaterally-biased hubs (highlighted in Figure 2D,E) likely play specialized roles in interhemispheric communication. In summary, the network’s topology is characterized by predominantly ipsilateral, intra-regional connections, with a minority of nodes mediating most long-range and cross-hemisphere communication.

### Spatial Dependence of Connectivity Strength

Given that the AMBCA provides standardized 3D coordinates for each brain region, we next examined how **physical distance** relates to connectivity strength. There is a clear spatial dependence in the mesoscale connectome: in general, brain regions that are nearer to each other tend to have denser connections, whereas weaker projections usually connect distant regions. This inverse relationship between Euclidean distance and projection density is visualized in Supplemental Figure 2, which plots average connection density against inter-region distance. Most high-density connections link regions that are anatomically close, reflecting the fact that many neural projections are localized. Meanwhile, connections bridging long distances (e.g., between forebrain and hindbrain structures) typically show lower density. Nevertheless, a few notable exceptions exist – cases where strong projections span large anatomical distances, suggesting specialized long-range communication channels.

To quantify these exceptions, we introduced a **weighted density** metric that combines connection strength with distance (Figure 2G). We multiplied each connection’s projection density by the Euclidean distance between source and target regions. This metric assigns greater weight to long-range connections that maintain high density. Using weighted density, we identified several pairs of brain regions that, despite being far apart, are linked by robust projections (appearing as high weighted-density outliers in Figure 2H). When averaging connectivity at the level of large brain divisions, we found that **intra-region connectivity not only dominates in strength but also tends to cover shorter physical distances**. For example, the cerebral nuclei and midbrain divisions have very high within-division connectivity (Figure 2F) and, correspondingly, relatively short average distances among their constituent nodes (Figure 2G). In contrast, inter-region connections often must span larger distances and generally have lower densities. However, a small number of long-distance links contribute significantly to the network’s integrated structure (as captured by the weighted density analysis). In summary, the strength of connections in the mouse brain has a strong spatial component: **most information travels along short-range, within-region pathways**. At the same time, a limited set of long-range projections provide critical bridges across distant parts of the brain.

### Higher-Order Network Properties

To understand the **network’s efficiency and integration** beyond direct connections, we analyzed higher-order connectivity measures such as the Degree of Separation (DOS; Figure 4) and weighted shortest paths (Figure 5). We first binarized the network at various density thresholds and computed the DOS between all pairs of nodes (i.e., the minimum number of synaptic steps required to connect one region to another). Remarkably, the mouse connectome exhibits very short path lengths, indicative of a **small-world organization**. Even after removing 75% of the weakest connections (retaining only edges with projection density > 0.75), the maximum DOS between any two brain areas was four. In other words, under a stringent threshold that preserves only the top quarter of connections, no region was more than four projections away from any other region. In the full, unfiltered network, most pairs of nodes are separated by only 2 or 3 steps (consistent with ∼92% edge density noted above), and the **network diameter** (longest shortest path) is effectively three (see Supplemental Figure 4A). This indicates a highly **high connectivity efficiency** – there are multiple redundant pathways such that information can travel from any source to target through just a few intermediate regions.

**Figure 4.**
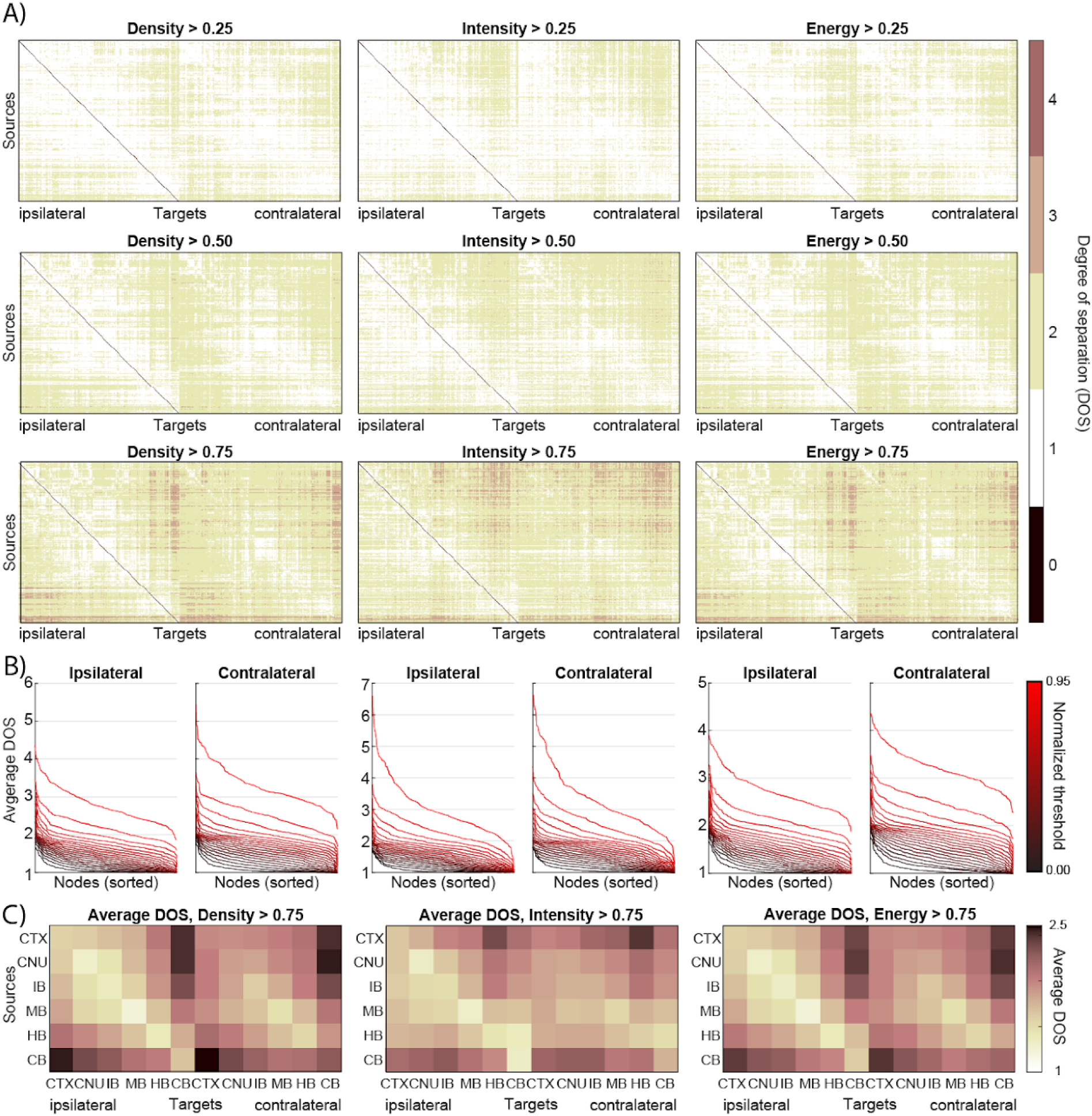
Degree of Separation (DOS) of the mouse brain network. **A)** Degree of Separation matrices for varying threshold and projection measure. In each matrix, an element (i,j) indicates the minimum number of edges necessary to walk from node i to node j. **B)** Average DOS from one source node to all others, shown for every node and sorted. Each line represents a threshold value, removing a different number of edges from the network. Data is shown for ipsi-and contralateral DOS and varying projection measures. **C)** The DOS matrix of every projection measure with Threshold=0.75 is condensed by averaging the nodes per larger brain area.

**Figure 5.**
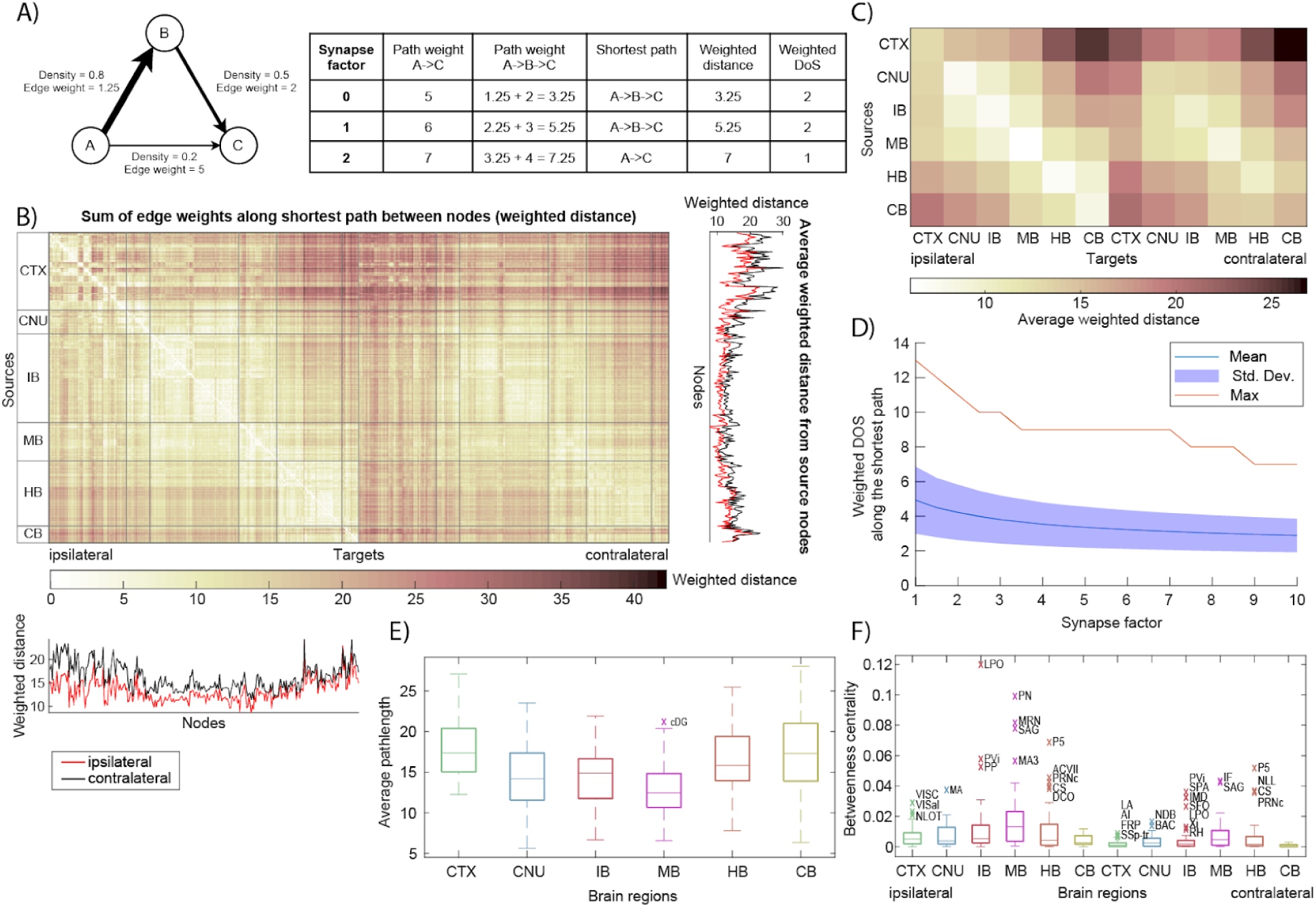
Shortest paths and weighted distance. **A)** Simple example network to clarify network distance measures. See the main text for explicit definitions. **B)** Matrix showing the weighted distance from every source node to every (bilateral) target node. Weighted distance is computed from the edge weight (1/projection density) using Dijkstra’s algorithm. Line plots on the bottom and right show the total input and output (i.e. sum over columns resp. rows of the adjacency matrix) per node separately for ipsi- (red) and contralateral (black) projections. **C)** Weighted distance matrix condensed by averaging over nodes per larger brain region. **D)** Weighted DOS is the number of edges along the shortest path between two nodes, the shortest path being the network path with the minimum weighted distance. The plot shows the weighted DOS as a function of synapse factor, a term added to edge weight to add extra distance for crossing synapses. **E)** For each larger brain region, the distribution of the average weighted distance to nodes from other brain regions is shown. Outliers are denoted with their node acronyms taken from AMBCA (see supplemental table 1). **F)** Betweenness centrality distributions per larger brain region, computed from the shortest path between every sorted pair of nodes.

As expected, increasing the threshold (thus pruning more connections) gradually increases path lengths; Figure 4B shows that the average DOS per node rises as the minimum edge density requirement is raised to 0.95. However, even at this very high threshold (keeping only the top 5% strongest connections), the network remains relatively well-connected, with most regions still reachable within a handful of steps. Furthermore, DOS analysis confirmed the earlier observation of autoconnectivity: when DOS matrices were averaged wi-thin each major brain region, within-region travel required the fewest steps (lowest DOS along the matrix diagonal), reflecting especially tight integration among subdivisions of the same region. Together, these results demonstrate a small-world topology in the mesoscale connectome – a dense core of connections ensures **short path lengths** and robust connectivity even when weaker links are ignored.

We next examined **weighted shortest paths** (Figure 5; Supplemental Figure 5) to incorporate connection strength into our assessment of network communication efficiency. Rather than treating all existing edges equally (as with DOS), we assigned a length to each connection based on its weight, using the inverse of projection density as the edge distance (so that stronger projections correspond to “shorter” distances). We then computed the minimal **weighted distance** between every pair of nodes using Dijkstra’s algorithm. The resulting weighted distance matrix (Figure 5B) provides a more nuanced view of network organization, highlighting how easily signals could travel between regions when favoring high-density pathways. From this analysis, we found that certain brain structures serve as particularly efficient bridges. Notably, the **Midbrain** (which here includes midbrain regions such as the thalamus and hypothalamus in the broader sense) has the **smallest average weighted distance** to all others. In other words, midbrain areas are, on average, only a short weighted distance away from any other part of the brain, underscoring their central integrative role in the connectome. By contrast, other divisions (such as the cerebellum or olfactory areas) remain more peripheral in terms of weighted distance, likely due to fewer or weaker long-range connections linking them to the rest of the brain.

We also assessed network **betweenness centrality** (BC) to identify potential hubs in information flow. BC was calculated for each node based on the fraction of all shortest paths (in the weighted network) that pass through that node. The distribution of BC values across regions revealed that most brain areas have low betweenness (many alternative routes exist), but a few standout nodes act as key intermediaries. For instance, the **Lateral Preoptic Area (LPO)** showed a very high BC (∼0.12), meaning about 12% of all shortest paths in the network go through this region. Such a value is an order of magnitude above the network average, highlighting LPO as an important hub (or bottleneck) for inter-regional communication. Other high-BC nodes (appearing as outliers in Figure 5F) similarly indicate brain areas that are disproportionately central for maintaining overall connectivity efficiency. These might correspond to major relay centers or integrative junctions in the brain. In summary, the analysis of higher-order properties confirms that the mouse brain network is **highly efficient and resilient**: most regions are only a few steps apart through either direct or indirect routes, and weighted-path analysis pinpoints specific regions that act as crucial connective hubs ensuring efficient signal propagation across the whole brain.

### Seeded analysis of networks in the brain

NeuroCarta could be used to explore specific subnetworks in the brain either by selective filtering of the input dataset, e.g., based on meta-variables like the sex (Figure 6), or identifying monosynaptically coupled networks that originate from a chosen set of structures, as in the sensorimotor connectivity circuit of the whisker system (Figure 7; (Heckman et al., 2017; Rault et al., 2024)).

**Figure 6.**
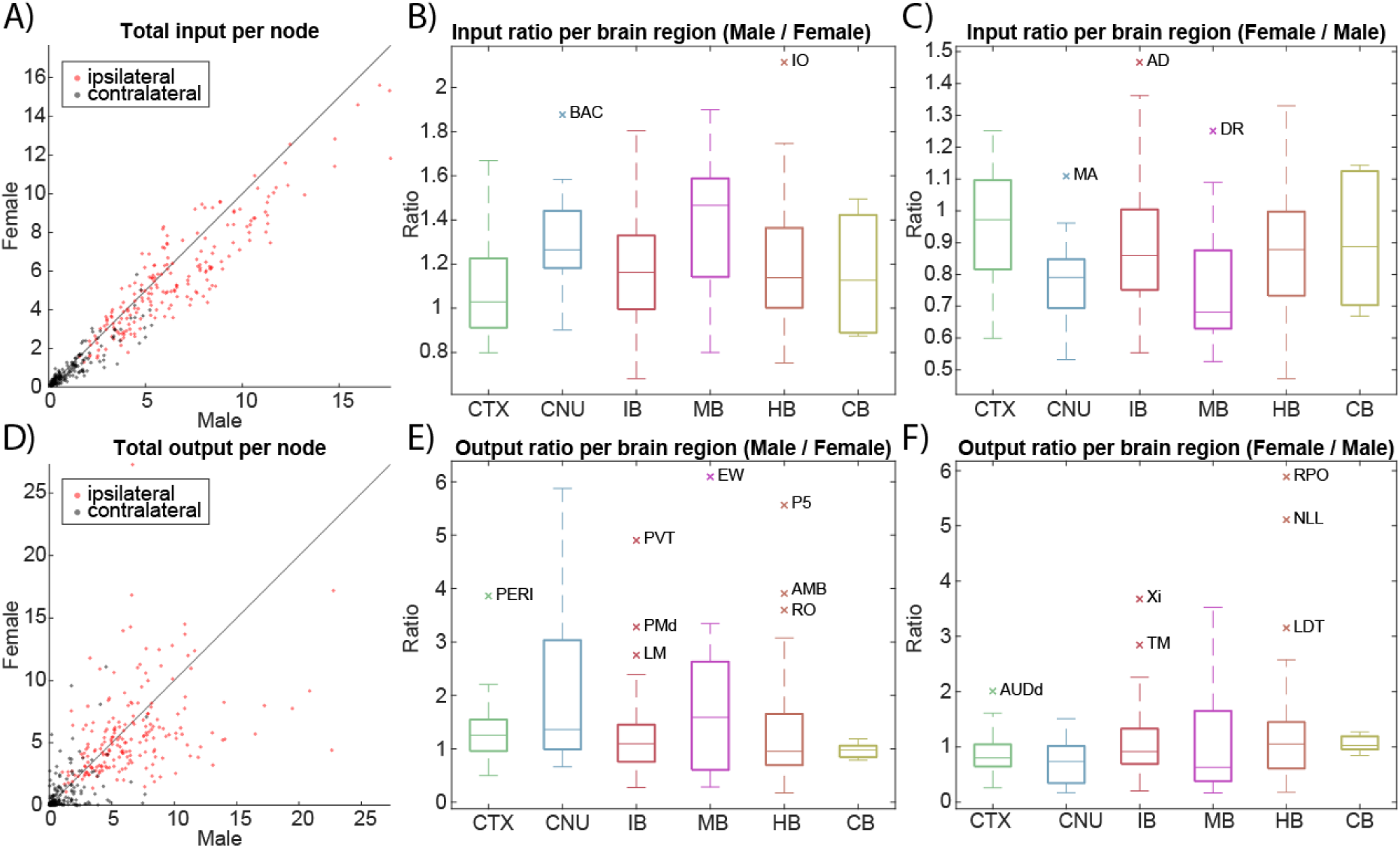
Sex-specific differences in connectivity. **A)** Total input per node shown for the male vs female mouse brain. **B)** The difference in node input in the male vs. female network is shown per larger brain area. Outliers are denoted with their node acronyms taken from AMBCA (see Supplemental table 1). **C)** The difference in node input in the female vs. male network is shown per larger brain area. **D)** Total output per node is shown for the male vs. female mouse brain. **E)** The difference in node output in the male vs. female network is shown per larger brain area. **F)** The difference in node output in the female vs. male network is shown per larger brain area.

**Figure 7.**
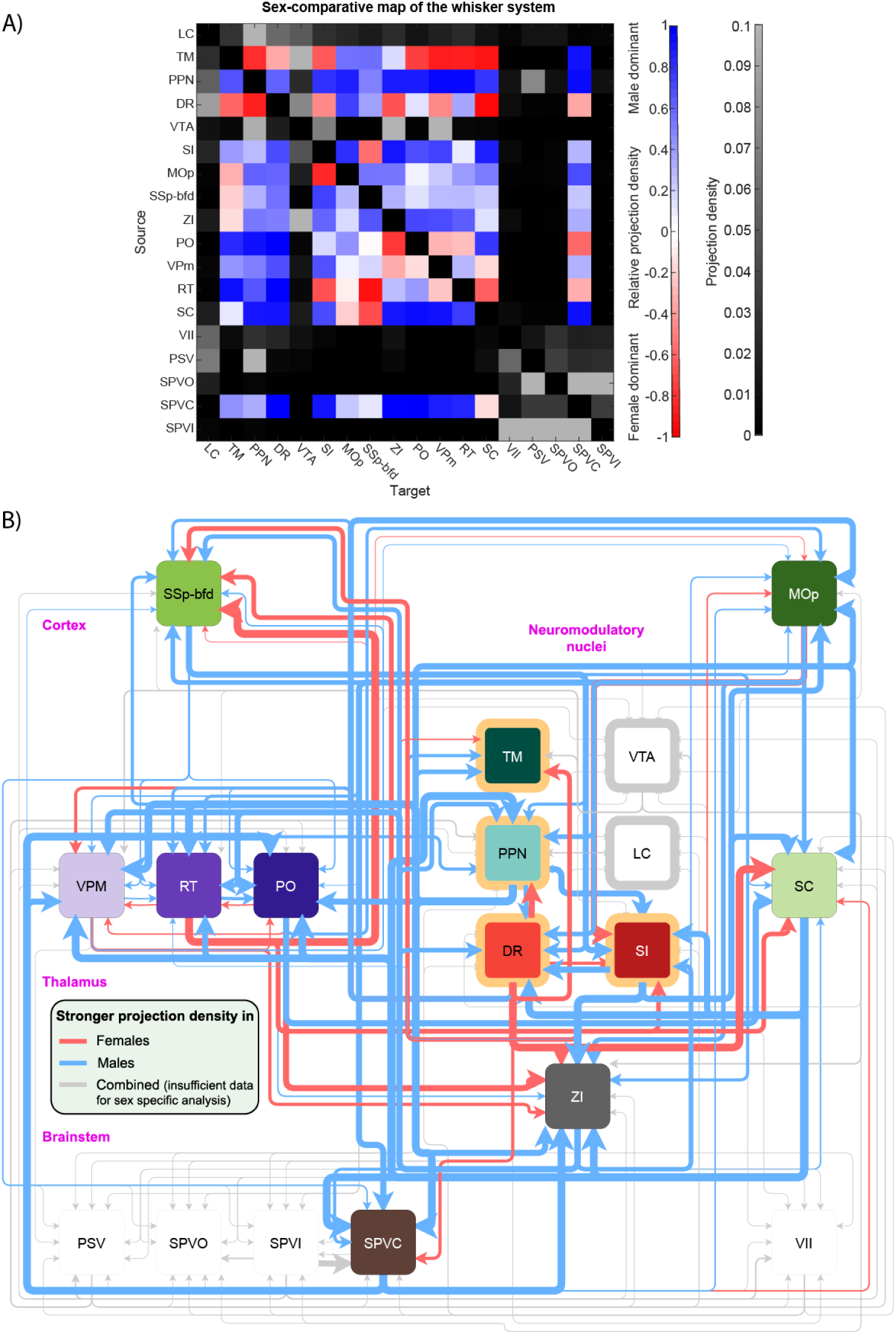
Sex-specific differences of connectivity in the whisker system. **A)** Adjacency matrix of the sex-comparative map of the whisker system. Colored matrix elements indicate stronger connections in male (blue) or female (red) mice. Grayscale elements indicate connections for which no sex-specific data is available and instead show regular projection density. Node acronyms are taken from AMBCA (see Supplemental Table 1). **B)** Network representation of the sex-comparative map of the whisker system. Edge color represents sex-specific overconnectivity in male (blue) or female (red) mice, and edge thickness indicates the quantity. Gray edges are existing connections for which no sex-specific data is available, edge thickness indicates regular projection density on a different scale.

We explored **sex-specific network differences** by constructing separate connectome models for male and female mice. The AMBCA data includes thousands of tracing experiments with recorded animal sex (1,758 male and 1,159 female in our dataset). We filtered the data to build an all-male network and an all-female network, each based on the subset of experiments conducted in mice of that sex. The initial male-derived network contained 255 node,s and the female network 218 nodes (since some brain regions had no data in one sex). After removing any region nodes that were not present in both, we obtained two comparable networks of 197 common nodes each for direct comparison. We then examined differences in the **total input and output connectivity profiles** for each region between males and females (Figure 6; Supplemental Figure 6).

Overall, the male and female connectivity matrices were highly correlated, but a number of regions showed significant quantitative differences in their connectivity strength. In Figure 6A and 6D, which plot total inputs and outputs per node for male vs. female, most points lie near the diagonal (indicating similar values in both sexes). However, the presence of several **outliers** reveals regions with notably different connectivity magnitudes. For instance, the **Inferior Olivary Complex** (a brainstem structure) receives more than **twice the total input** in male mice compared to females. Conversely, the **Anterodorsal Thalamic Nucleus** receives about 1.5× greater input in females than males. These differences suggest sex-specific variation in how strongly certain areas are targeted by incoming projections.

On the output side, we found a few regions with even more dramatic disparities: the **Edinger–Westphal nucleus** (a midbrain nucleus) projects roughly **6 times more strongly in males** than in females, whereas the **Nucleus raphe pontis** (a hindbrain area) projects much more strongly in females (up to six-fold difference). These data are summarized in Figure 6B,C (for inputs) and Figure 6E,F (for outputs), which show the distribution of male-vs-female differences by major region, with the aforementioned regions marked as outliers. Such node-level differences imply that certain circuits (for example, those involving the oculomotor system, of which Edinger–Westphal is a part, or the arousal pathways via raphe nuclei) may be wired with different strengths in male versus female brains.

We created a comparative connectivity map of a well-characterized network: the mouse whisker (somatosensory) system to visualize where these sex-biased differences occur in a circuit context. This system involves a series of projections linking the whisker follicles to the cortex through the brainstem and thalamic relays (as described in (Rault et al., 2024)). We extracted the corresponding subgraph from our male and female networks using a set of key regions and connections identified for the whisker system in prior work. We then computed the **relative projection density** for each connection in this subgraph, which is defined as the ratio or difference between the male and female projection strengths (see Methods). The resulting **sex-comparative adjacency matrix** is shown in Figure 7A, and a weighted, directed network graph is shown in Figure 7B. The corresponding projection density-based adjacency matrix is shown in Supplemental Figure 7.

In summary, the **sex-specific analysis** showed that the overall mesoscale connectome is largely similar in male and female mice but with **specific quantitative differences** in connectivity that stand out in certain regions and pathways. These differences were detectable both at the global level (total inputs/outputs of certain nodes differ by more than two-fold between sexes) and at the circuit level (particular sensory pathways have biased connection strengths). Such findings suggest potential anatomical bases for sex differences in sensory processing or other behaviors, and they demonstrate how NeuroCarta can be used to uncover fine-grained network differences in subset populations. Altogether, the Results illustrate the versatility of the NeuroCarta toolbox in analyzing the mouse brain connectome — from global network structure and spatial organization to predictive modeling, as well as parsing the connectome by cell type and sex to reveal biologically meaningful variations in connectivity.

## Discussion

NeuroCarta is a computational toolbox that leverages the AMBCA dataset for automated, large-scale network construction and analysis. By integrating anatomical connectivity data into a quantitative network framework, NeuroCarta enables researchers to extract insights into brain connectivity topology, interregional communication, and global network efficiency. The toolbox facilitates the conversion of raw connectivity data into weighted and directed graphs, allowing users to systematically investigate properties such as degree of separation, connectivity strength, clustering, and centrality measures. Given the growing reliance on computational approaches in connectomics, NeuroCarta provides an essential tool for examining how mesoscale connectivity shapes neural processing and functional interactions.

## Limitations and Considerations

While NeuroCarta is a powerful tool for mesoscale connectivity analysis, several inherent limitations should be considered when interpreting results.

The accuracy and completeness of the networks constructed using NeuroCarta directly depend on the experimental scope of the AMBCA. As noted previously (Smith et al., 2024), the segmentation accuracy and anatomical resolution of connectivity mappings can be affected by the number of viral tracer injections and the algorithmic approaches used in data processing. The NeuroCarta incorporates thresholding and filtering methods to improve signal-to-noise ratio, but future refinements could benefit from additional cross-validation with independent datasets.

As a tool focused on anatomical network construction, NeuroCarta does not incorporate synaptic weights, neuronal activity levels, or functional interactions between regions. While the network-based approach in NeuroCarta enables the calculation of degree of separation, clustering coefficients, and centrality measures, these metrics are context-dependent and should not be overinterpreted without functional validation. Specific attractor nodes identified in the network may be anatomical hubs, for example, but their involvement in information processing pathways requires additional physiological validation. Although anatomical connectivity provides a foundation for functional network modeling, future work integrating fMRI, calcium imaging, or optogenetic data could bridge this gap and enable comparative structure-function analyses.

The toolbox operates at a mesoscale resolution, where nodes correspond to brain regions rather than individual neurons or microcircuits. While this approach allows for efficient whole-brain analyses, it does not capture fine-grained synaptic specificity or neuronal subtype connectivity. Researchers interested in circuit-level interactions may need to complement NeuroCarta with single-cell resolution tracing datasets or electrophysiological recordings.

## Applications

Despite data-related limitations, NeuroCarta provides a powerful and versatile framework for studying mesoscale brain connectivity. One of its key applications is comparative network analysis, e.g., sex-specific differences in connectivity as quantified in this study. Beyond available metavariables, e.g., sex differences, transgenic lines, and mouse strain, the toolbox can import independent data to explore developmental changes, genetic influences, and disease-associated alterations in neural connectivity. The flexibility of NeuroCarta allows for the customization of network construction, facilitating research on specific circuit modules, neurotransmitter-defined pathways, or large-scale anatomical variations across the brain.

Another significant application of NeuroCarta is in neuroinformatics. The connectivity matrices generated by the toolbox can be exported to external graph theory toolboxes, such as the Brain Connectivity Toolbox (BCT) (Rubinov & Sporns, 2010), and integrated into neural network simulations. In future studies, the toolbox could be expanded to include machine learning algorithms, allowing researchers to predict missing connections, identify recurrent network motifs, and classify connectivity patterns under different experimental conditions.

Another major advantage of NeuroCarta is its ability to generate testable hypotheses about neural circuit function. By quantifying anatomical network properties, the toolbox can guide hypothesis-driven experimental research. For example, if a particular brain region emerges as a high-degree hub in network analysis, optogenetics, calcium imaging, electrophysiology and behavioral analysis can be employed or multimodal datasets could be utilized, see e.g. the whisker system (Azarfar, Zhang, et al., 2018; da Silva Lantyer et al., 2018; Kole, Komuro, et al., 2017; Kole, Lindeboom, et al., 2017), to examine its role in sensorimotor integration or cognitive processing. This structure-function approach provides an iterative framework in which computational network models inform experimental design, leading to new insights into brain organization.

Beyond rodent studies, NeuroCarta can be extended to cross-species comparisons. Although the toolbox is currently optimized for mouse brain connectivity, its workflow could be adapted to analyze anatomical tracing data from other species, including non-human primates and humans. Incorporating human diffusion MRI data or non-human primate connectomes into the analysis could enhance our understanding of evolutionary differences in brain organization. Such comparative studies could provide insights into species-specific adaptations in network structure and function, offering a broader perspective on brain evolution and cognition.

By integrating quantitative network analysis with experimental neuroscience, computational modeling, and translational applications, NeuroCarta serves as an essential tool for advancing connectomics research. Its adaptability across multiple domains ensures that it will continue to play a pivotal role in mapping, analyzing, and interpreting neural networks in both health and disease.

## Conclusion

The increasing availability of large-scale anatomical datasets presents new opportunities for quantifying and analyzing brain connectivity, but also introduces challenges in data integration, processing, and interpretation. NeuroCarta provides a scalable, user-friendly solution for constructing and analyzing mesoscale connectivity networks, bridging the gap between raw anatomical data and network-based neuroscience. By automating connectivity quantification, facilitating graph-theoretic analyses, and enabling cross-modality integration, NeuroCarta serves as a quantitative platform to investigate fundamental principles of brain organization, network topology, and neural computation. Future extensions of NeuroCarta will focus on multi-modal integration with gene expression datasets, functional imaging data, and advanced predictive modeling, further enhancing its potential as a comprehensive framework for connectomics and network neuroscience research.

## Supplemental figures and tables

**Supplemental figure 1.**
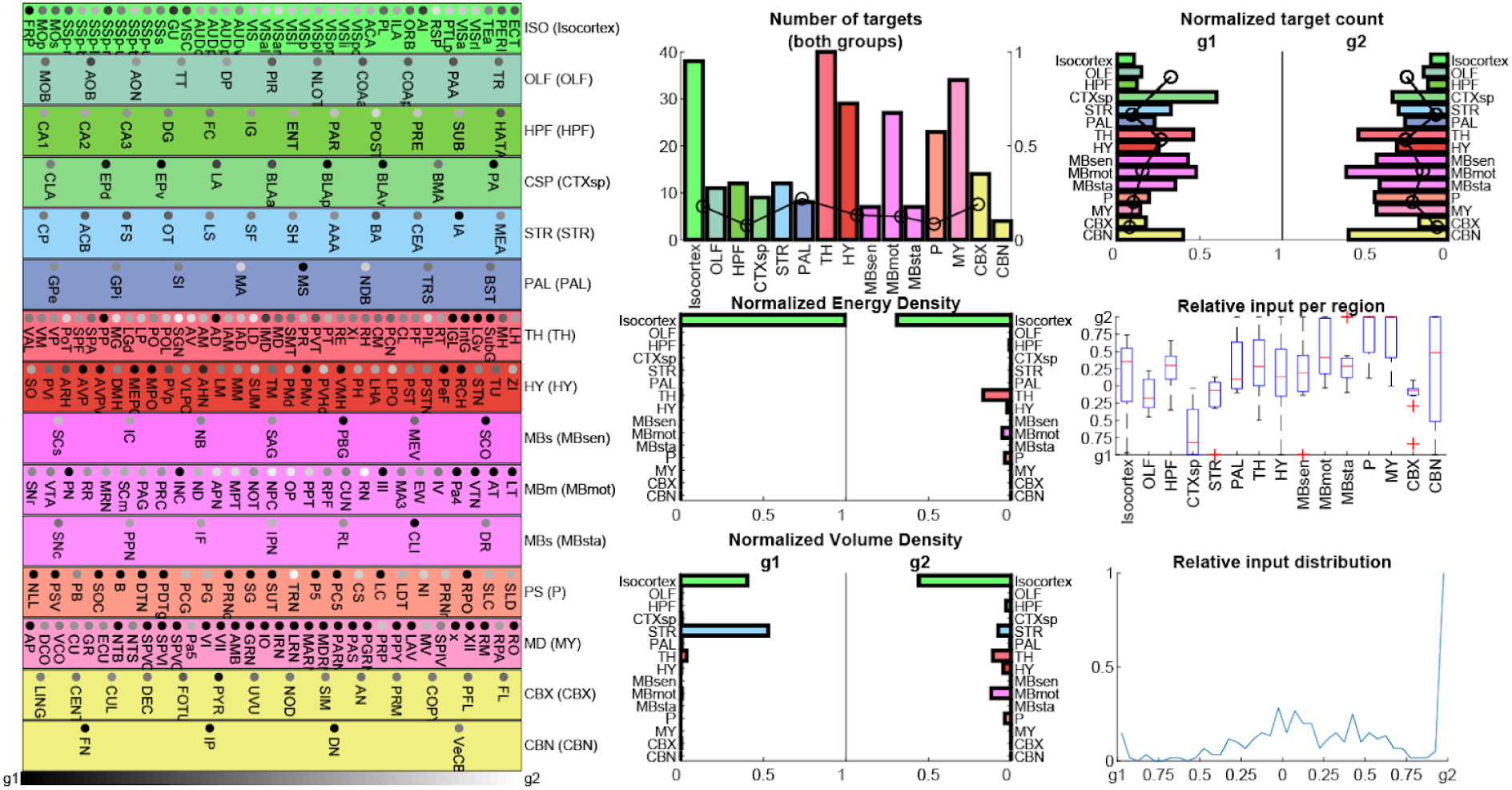
Example of the function output of the *crosscorrelatemaps* function. The *crosscorrelatemaps* function generates a MATLAB figure showing statistical differences between two experiments. Specifically, this example compares the experiments with ids 100140756 (indicated in the figure as g1) and 100140949 (indicated as g2). The figure on the left shows the relative projection density of projections towards postsynaptic targets, originating in the experiments’ respective projection sites. The smaller figures on the right show various statistical distributions comparing the two experiments, i.e. the total number of postsynaptic targets in the two experiments combined; the normalized number of targets, projection density, and projection volume per larger brain area shown separately for the two experiments; and the distributions of incoming projections ratios between the two experiments.

**Supplemental figure 2.**
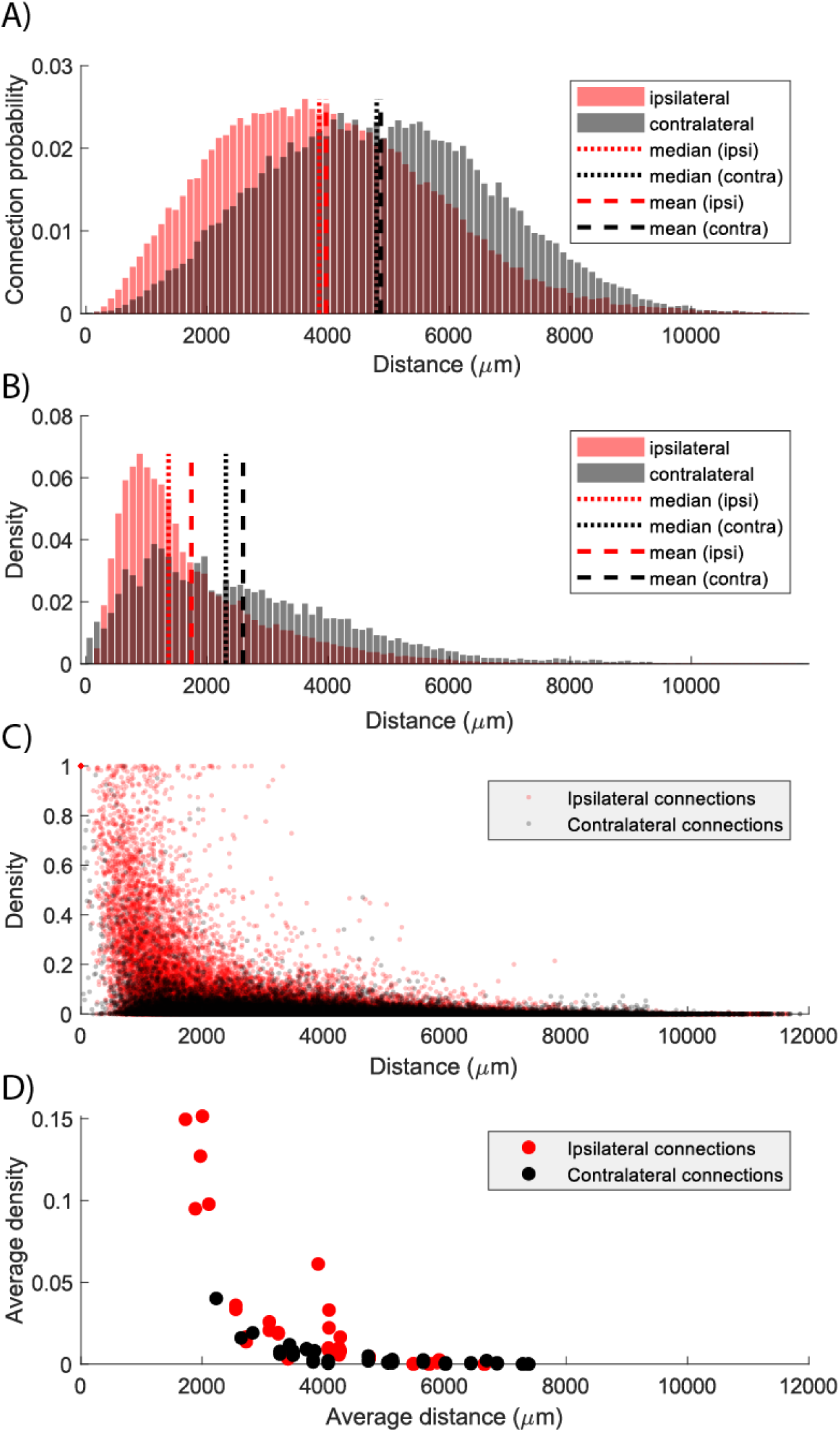
Projection density declines over growing euclidean distance. **A)** The probability of connection (i.e. number of connections normalized to the area under curve) as a function of Euclidean distance between the presynaptic source of the projection and postsynaptic target, shown separately for ipsi- and contralateral projections. Distances were calculated from the geometric centers of the targets. **B)** Distribution of projection densities as a function of Euclidean distance. **C)** Projection density plotted against Euclidean distance for all existing ipsi- and contralateral projections. **D)** Projection density plotted against Euclidean distance, averaged per larger brain region.

**Supplemental figure 3.**
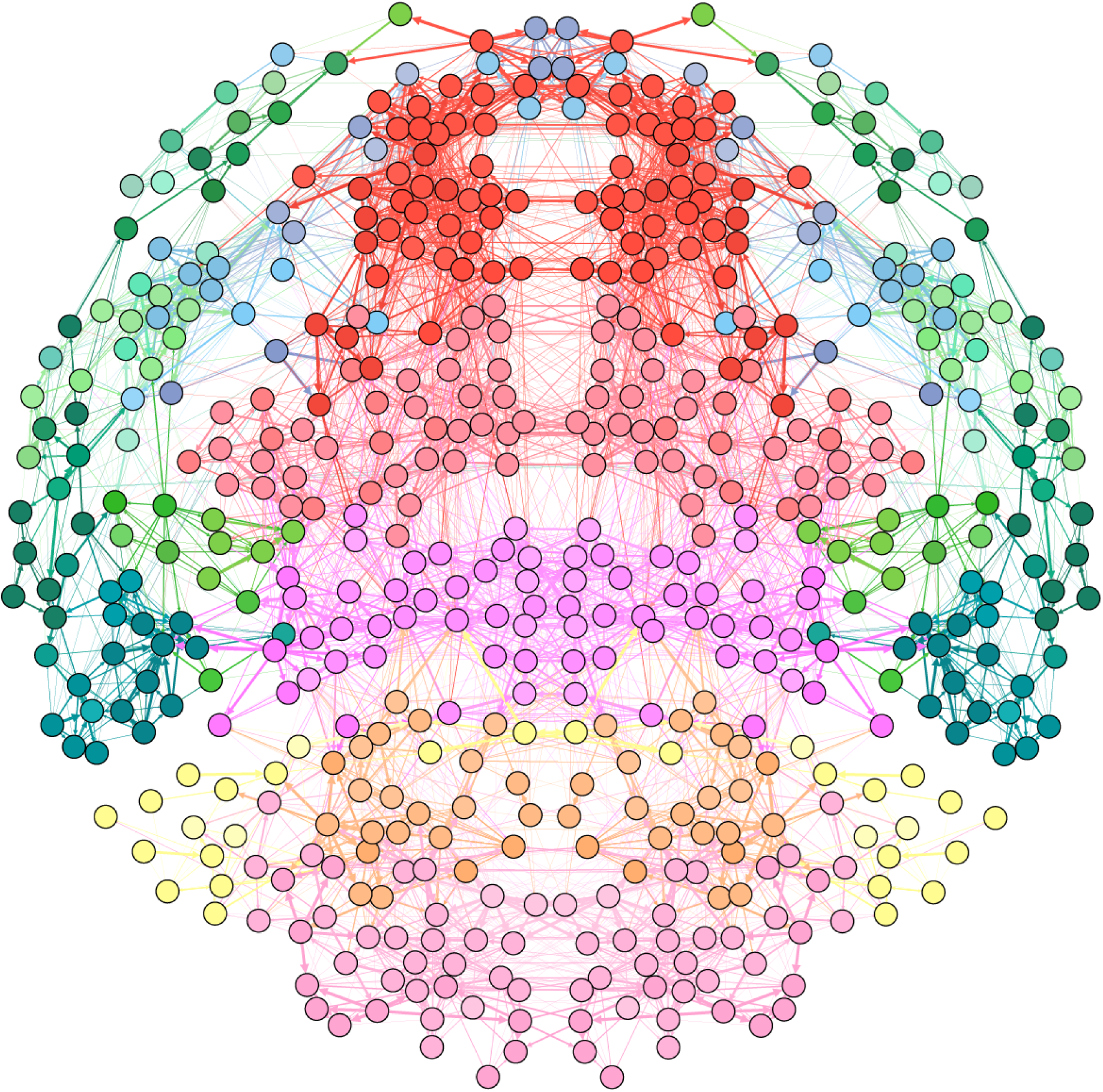
Bilateral mouse brain visualized in Gephi. The density-based mouse brain network constructed by Neurocarta was exported in GEXF file format and then visualized in Gephi. Using the Fruchterman-Reingold algorithm, a network layout was generated based solely on the network connectivity. For visibility reasons, only 10% of the strongest edges are shown. Node coloring is the same as used in the AMBCA (green = cortex, red = interbrain, purple = midbrain and hindbrain, yellow = cerebellum).

**Supplemental figure 4.**
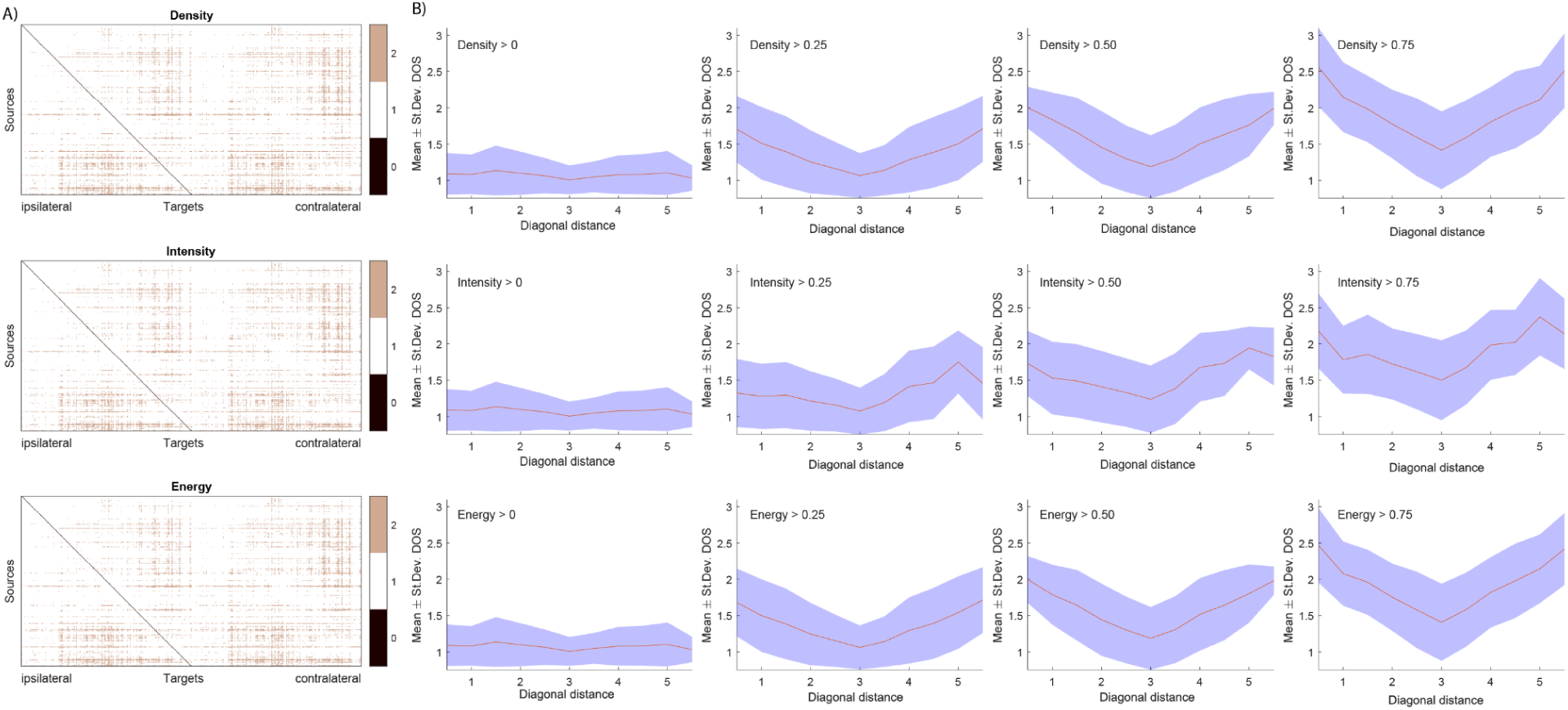
Degree of separation. **A)** The degree of separation (DOS) matrices for an non-thresholded bilateral, density-based shown for networks based on projection density, intensity and energy. **B)** The average DOS across the positive diagonals of the respective averaged DOS matrices from Figure 4A, showing that DOS is generally lower within the larger brain regions.

**Supplemental figure 5.**
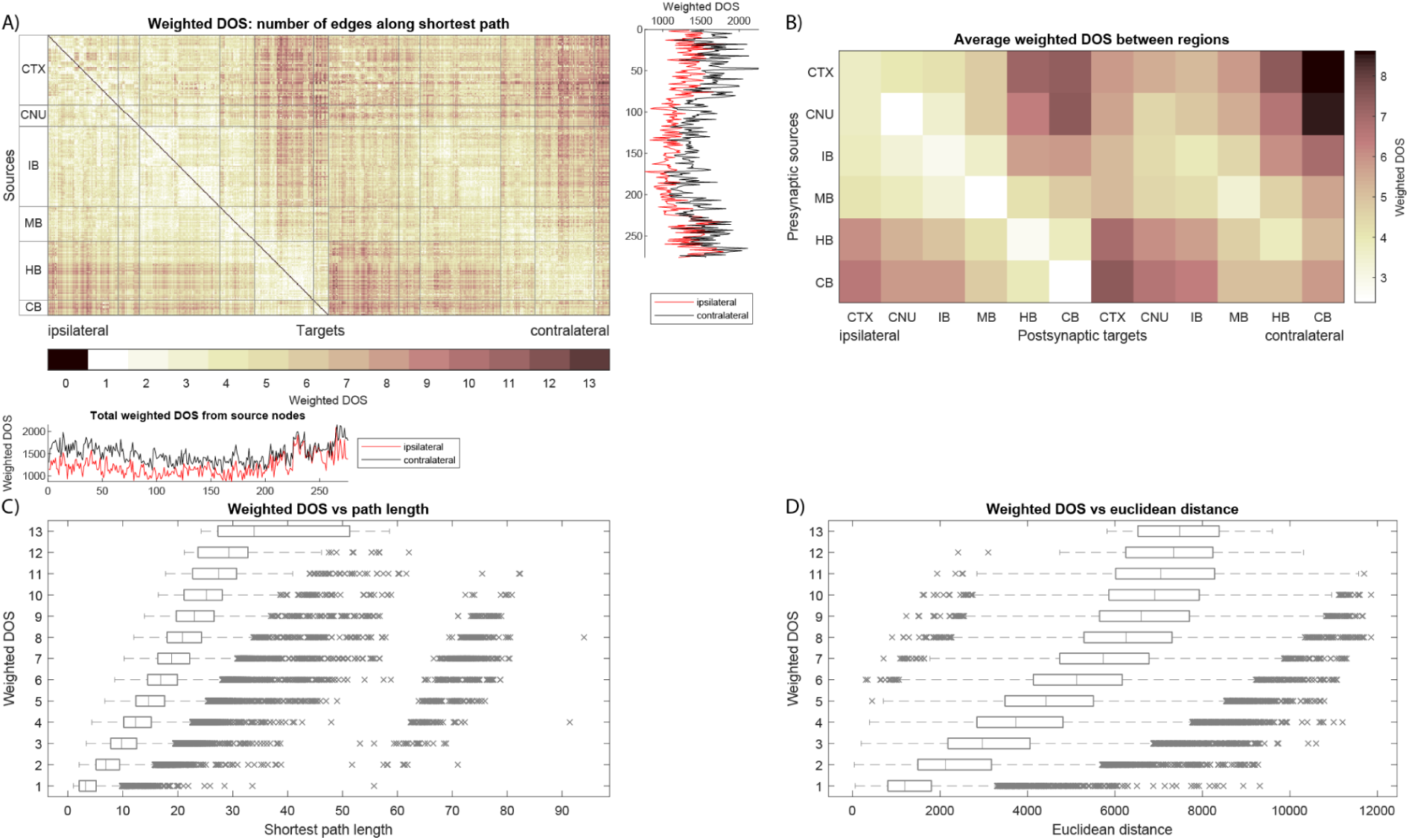
Weighted degree of separation. **A)** Weighted degree of separation (weighted DOS) between any pair of nodes in the bilateral, density-based network. Row-wise sums (for outgoing weighted DOS) and column-wise sums (for incoming weighted DOS) are shown in the line plots on the sides. **B)** Weighted DOS averaged within larger brain areas. **C)** Distribution of weighted distances for each occurring value of weighted DOS, showing the relation between the two. **D)** Distribution of euclidean distances for each occurring value of weighted DOS.

**Supplemental figure 6.**
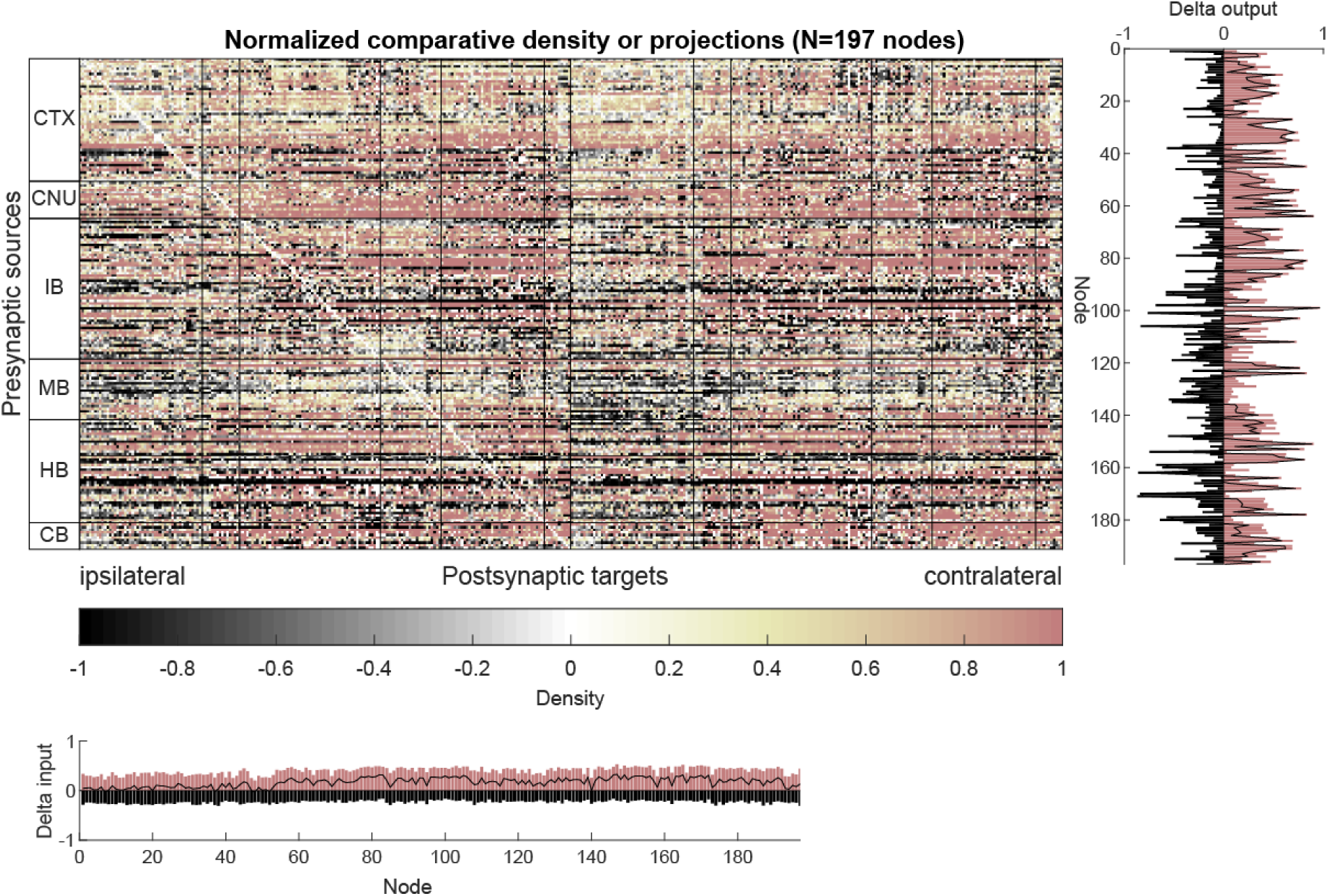
Comparative map between sexes. Comparative density shown for all nodes in the bilateral network. A value of -1 means a projection only exists in female mice, a value of 1 means a projection only exists in male mice, a value of 0 means the connection has equal strength in both sexes, and any value in between indicates a connection that is relatively stronger in one of the sexes. The area plots on the sides show the row-wise and column-wise sums of the matrix, indicating which brain areas either receive more incoming projections in a particular sex, or send more outgoing projections.

**Supplemental figure 7.**
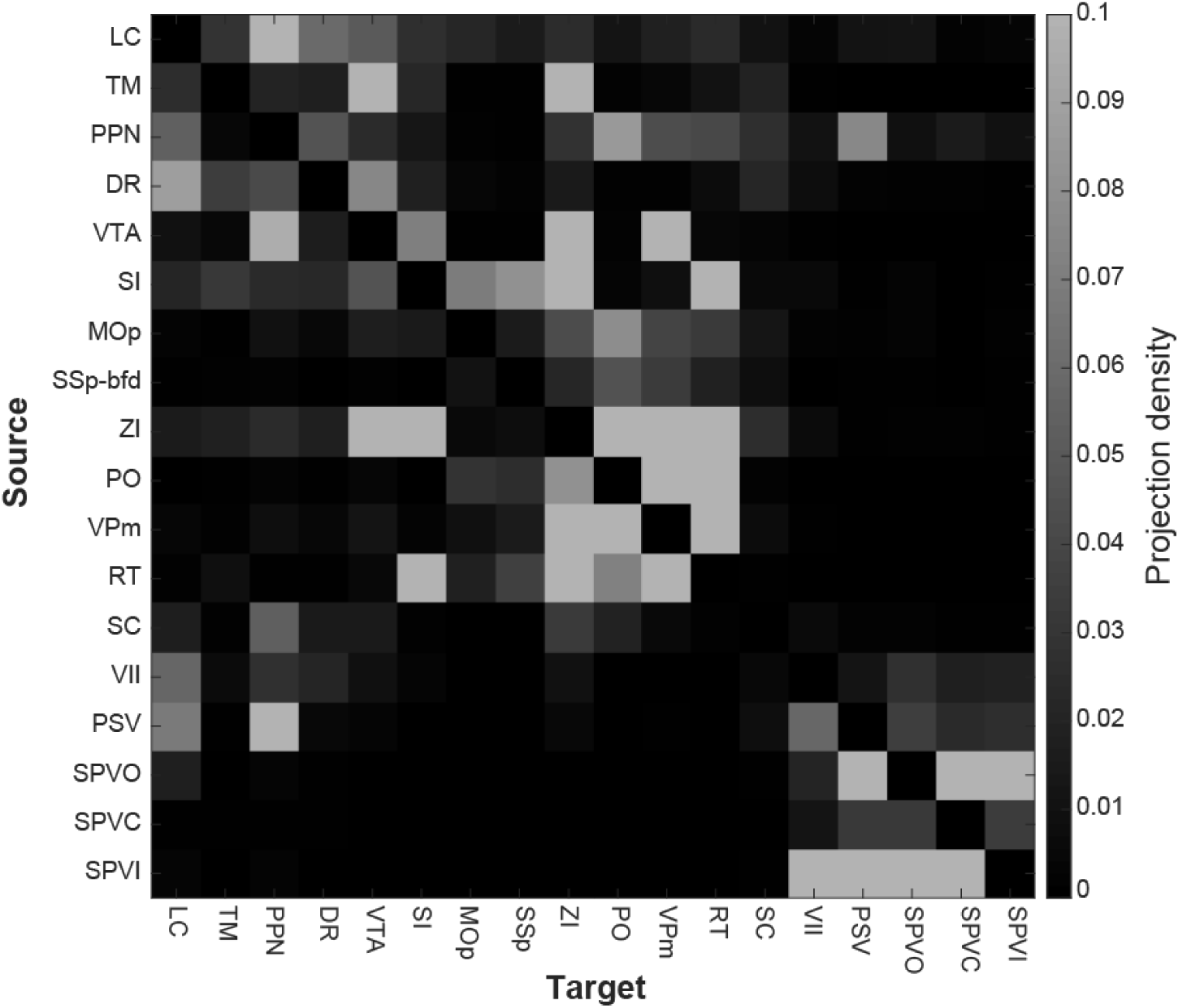
Connectivity matrix of the whisker system. Density-based connectivity matrix of the whisker system using the nodes selected by Raoult et al. (Raoult et al. 2024). This matrix contains the same data as the one in Figure 7A except for omitting the sex-specific projections.

**Supplemental table 1:** Node acronyms.

| **Acronym** | **Brain structure name (taken from AMBCA)** |
| --- | --- |
| ACVII | Accessory facial motor nucleus |
| AD | Anterodorsal nucleus |
| AI | Agranular insular area |
| AMB | Nucleus ambiguus |
| AUDd | Dorsal auditory area |
| BAC | Bed nucleus of the anterior commissure |
| CB | Cerebellum |
| CBN | Cerebellar nuclei |
| CBX | Cerebellar cortex |
| cDG | Dentate gyrus (contralateral to source node) |
| CM | Central medial nucleus of the thalamus |
| CNU | Cerebral nuclei |
| COPY | Copula pyramidis |
| CP | Caudoputamen |
| CS | Superior nucleus raphe |
| CTX | Cerebral cortex |
| CTXsp | Cortical subplate |
| DCO | Dorsal cochlear nucleus |
| DG | Dentate gyrus |
| DR | Dorsal nucleus raphe |
| EW | Edinger-Westphal nucleus |
| FRP | Frontal pole |
| HB | Hindbrain |
| HPF | Hippocampal formation |
| HY | Hypothalamus |
| IB | Interbrain |
| IF | Interfascicular nucleus raphe |
| ILA | Infralimbic area |
| IMD | Interomedial dorsal nucleus of the thalamus |
| IO | Inferior olivary complex |
| LA | Lateral amygdalar nucleus |
| LC | Locus coeruleus |
| LDT | Laterodorsal tegmental nucleus |
| LING | Lingula |
| LM | Lateral mammillary nucleus |
| LPO | Lateral preoptic area |
| MA | Magnocellular nucleus |
| MA3 | Medial accessory oculomotor nucleus |
| MB | Midbrain |
| MBmot | Midbrain, motor related |
| MBsen | Midbrain, sensory related |
| MBsta | Midbrain, behavioral state related |
| MEPO | Median preoptic nucleus |
| MOp | Primary motor area |
| MRN | Midbrain reticular nucleus |
| MS | Medial septal nucleus |
| MY | Medulla |
| NDB | Diagonal band nucleus |
| NLL | Nucleus of the lateral lemniscus |
| NLOT | Nucleus of the lateral olfactory tract |
| OLF | Olfactory areas |
| ORB | Orbital area |
| P | Pons |
| P5 | Peritrigeminal zone |
| PAG | Periaqueductal gray |
| PAL | Pallidum |
| PERI | Perirhinal area |
| PL | Prelimbic area |
| PMd | Dorsal premammillary nucleus |
| PN | Paranigral nucleus |
| PO | Posterior complex of the thalamus |
| PP | Peripeduncular nucleus |
| PPN | Pedunculo pontine nucleus |
| PRNc | Pontine reticular nucleus, caudal part |
| PSV | Principal sensory nucleus of the trigeminal |
| PVi | Periventricular hypothalamic nucleus, intermediate part |
| PVT | Paraventricular nucleus of the thalamus |
| RH | Rhomboid nucleus |
| RO | Nucleus raphe obscurus |
| RPO | Nucleus raphe pontis |
| RSP | Retrosplenial area |
| RT | Reticular nucleus of the thalamus |
| SAG | Nucleus sagulum |
| SC | Superior colliculus |
| SFO | Subfornical organ |
| SI | Substantia innominata |
| SPA | Subparafascicular area |
| SPVC | Spinal nucleus of the trigeminal, caudal part |
| SPVI | Spinal nucleus of the trigeminal, interpolar part |
| SPVO | Spinal nucleus of the trigeminal, oral part |
| SSp-bfd | Primary somatosensory area, barrel field |
| SSp-tr | Primary somatosensory area, trunk |
| STR | Striatum |
| TH | Thalamus |
| TM | Tuberomammillary nucleus |
| TRS | Triangular nucleus of septum |
| VII | Facial motor nucleus |
| VIS | Visual areas |
| VISal | Anterolateral visual area |
| VISC | Visceral area |
| VISp | Primary visual area |
| VPM | Ventral posteromedial nucleus of the thalamus |
| VTA | Ventral tegmental area |
| Xi | Xiphoid thalamic nucleus |
| ZI | Zona incerta |

